# Differential screening identifies molecules specifically inhibiting CCR5 transport to the cell surface and HIV infection

**DOI:** 10.1101/364927

**Authors:** Gaelle Boncompain, Floriane Herit, Sarah Tessier, Aurianne Lescure, Elaine Del Nery, Pierre Gestraud, Isabelle Staropoli, Yuko Fukata, Masaki Fukata, Anne Brelot, Florence Niedergang, Franck Perez

## Abstract

Proteins destined to the cell surface are conveyed through membrane-bound compartments using the secretory pathway. Multiple secretory routes exist in cells, which paves the way to the development of inhibitory molecules able to specifically perturb the transport of a chosen cargo. We used differential high-content screening of chemical libraries to identify molecules reducing the secretion of CCR5, the major co-receptor for HIV-1 entry. Three molecules strongly affected the anterograde transport of CCR5, without inhibiting the transport of the related G protein-coupled receptors CCR1 and CXCR4. These three molecules perturb the transport of endogenous CCR5 and decrease the entry of HIV in human primary target cells. Two molecules were found to share the same mode of action, inhibiting palmitoylation of CCR5. Our results demonstrate that secretory routes can be specifically targeted which allows to envisage novel strategies to provoke the intracellular retention or rerouting of secretory proteins involved in disease development.

## Introduction

The Human Immunodeficiency Virus type 1 (HIV-1) infects immune cells, in particular, CD4^+^ T lymphocytes and macrophages, leading to acquired immunodeficiency syndrome (AIDS). The cell entry of HIV-1 is initiated by the interaction of its surface envelope glycoprotein, gp120, with two host cell surface receptors: CD4 and a co-receptor. The CC chemokine receptor 5 (CCR5) is the principal co-receptor for R5-tropic strains, responsible for the transmission and establishment of HIV-1 infection ^1-5^. Genetic polymorphism in the *CCR5* gene has been correlated with HIV resistance. Individuals homozygote for the CCR5 delta32 allele do not express CCR5 at the cell surface and are resistant to HIV-1 infection ^6, 7^. An additional demonstration of the crucial role CCR5 plays in HIV-1 infection came from the long-term control of infection in a patient transplanted with stem cells from a delta32/delta32 individual ^8^. Importantly, CCR5 delta32 individuals do not show major deficiencies due to the absence of cell surface CCR5 and as such, the therapy shows great promise.

Consequently, several anti-HIV therapies targeting CCR5 have been developed such as the drug Maraviroc ^9, 10^, CCR5 blocking antibodies ^11, 12^ and *CCR5* gene editing ^13, 14^. Of these, Maraviroc is the only anti-HIV therapy targeting CCR5 currently used for the treatment of patients. By binding to CCR5, this small, nonpeptidic CCR5 ligand prevents the interaction of the HIV-1 gp120 to CCR5 via an allosteric mechanism. Like the majority of therapies targeting CCR5, Maraviroc aims at inhibiting CCR5 interaction with HIV gp120 at the cell surface.. Our approach takes advantage of the diversity of the secretory routes ^15^ to selectively inhibit the secretion of CCR5 at the cell surface without perturbing the overall protein transport.

CCR5 is a member of the class A G-protein coupled receptor (GPCR) family, containing seven transmembrane domains which enters the secretory pathway at the level of the endoplasmic reticulum (ER). It is then exported from the ER to the Golgi complex and then to the cell surface. Little is known about the molecular motifs involved in the transport of CCR5 to the cell surface along the secretory pathway.

It has been recently revealed that a diversity of secretion routes, which have long been overlooked, exist for different cargos in mammalian cells ^15^. These observations allowed us to take advantage of this diversity to selectively inhibit the secretion of CCR5 at the cell surface without broadly perturbing protein transport. We used the Retention Using Selective Hooks (RUSH) assay ^16^ to synchronize the anterograde transport of CCR5 and allow quantitative monitoring of the protein’s transport. It enabled the high-content screening of chemical libraries to identify small molecules able to inhibit the secretion of CCR5 to the cell surface. We showed that the effects of two of these molecules depend on cysteine residues present in the cytoplasmic tail of CCR5. These cysteine residues are target of palmitoylation and we observed that the two molecules strongly reduced their modification. A significant reduction in HIV-1 entry and *de novo* virus production by target cells was observed after treatment with the identified molecules.

Together, our data indicate that perturbation of CCR5 modification, and more generally, that the diversity of secretory pathway represents an important and underexploited source for drug discovery.

## Results

### Differential transport of CCR5 and TNF to the cell surface

The quantity of CCR5 present at the cell surface at steady state corresponds to a balance between the secretion of newly synthetized CCR5 and its endocytosis, recycling and degradation. To study the secretion of CCR5 and identify compounds that affect its anterograde transport, its transport was synchronized using the Retention Using Selective Hooks (RUSH) assay ^16^. Briefly, the cargo of interest is fused to a Streptavidin Binding Peptide (SBP) and co-expressed with a resident protein of the endoplasmic reticulum (ER) which is fused to Streptavidin. The streptavidin ‘hook’ retains the cargo upon synthesis due to Streptavidin-SBP interaction and prevents its export from the ER. The synchronized transport of the cargo is induced by the addition of biotin that rapidly enters cells, binds to Streptavidin and competes out SBP. HeLa cells stably expressing GFP-tagged CCR5, adapted to the RUSH assay, were established (Str-KDEL_SBP-EGFP-CCR5). In the absence of biotin, CCR5 was localized in the endoplasmic reticulum (**Fig. 1a, 0 min**). Addition of biotin enabled export of CCR5 from the ER towards the Golgi complex and its subsequent appearance at the cell surface (**Fig. 1a, 30 min – 120 min.**). The transport of CCR5 to the cell surface was compared to the transport of another RUSH-adapted cargo, TNF ^16, 17^. While transport intermediates were visible from ER to Golgi and from Golgi to plasma membrane for TNF, very few were detected for CCR5 (**Fig. 1a, Supplementary videos 1 and 2**). The EGFP is exposed to the extracellular face of the plasma membrane in both CCR5 and TNF constructs. The amount of cargo arriving at the cell surface over time was assessed using flow cytometry of non-permeabilized cells labeled with an anti-GFP antibody. CCR5 was transported more slowly than TNF to the plasma membrane (**Fig. 1b**). 90 min after release from the ER, the amount of CCR5 at the cell surface was stable, while TNF rapidly disappeared from the cell surface. These results are consistent with our previous observations demonstrating that TNF is rapidly internalized from the surface of HeLa cells ^16^. In contrast, CCR5 was found to be stably expressed at the cell surface. Analysis of the simultaneous trafficking of co-expressed CCR5 and TNF confirmed the slower kinetics of CCR5 and revealed the segregation of the two cargos at the level of the Golgi complex. 20 min after release from the ER, TNF was exported from the Golgi complex in tubular and vesicular transport carriers from which CCR5 was excluded (**Fig. 1c, d, Supplementary videos 3**). Taken together, these results suggest that distinct molecular machineries are involved in the secretion of CCR5 and TNF.

**Figure 1:**
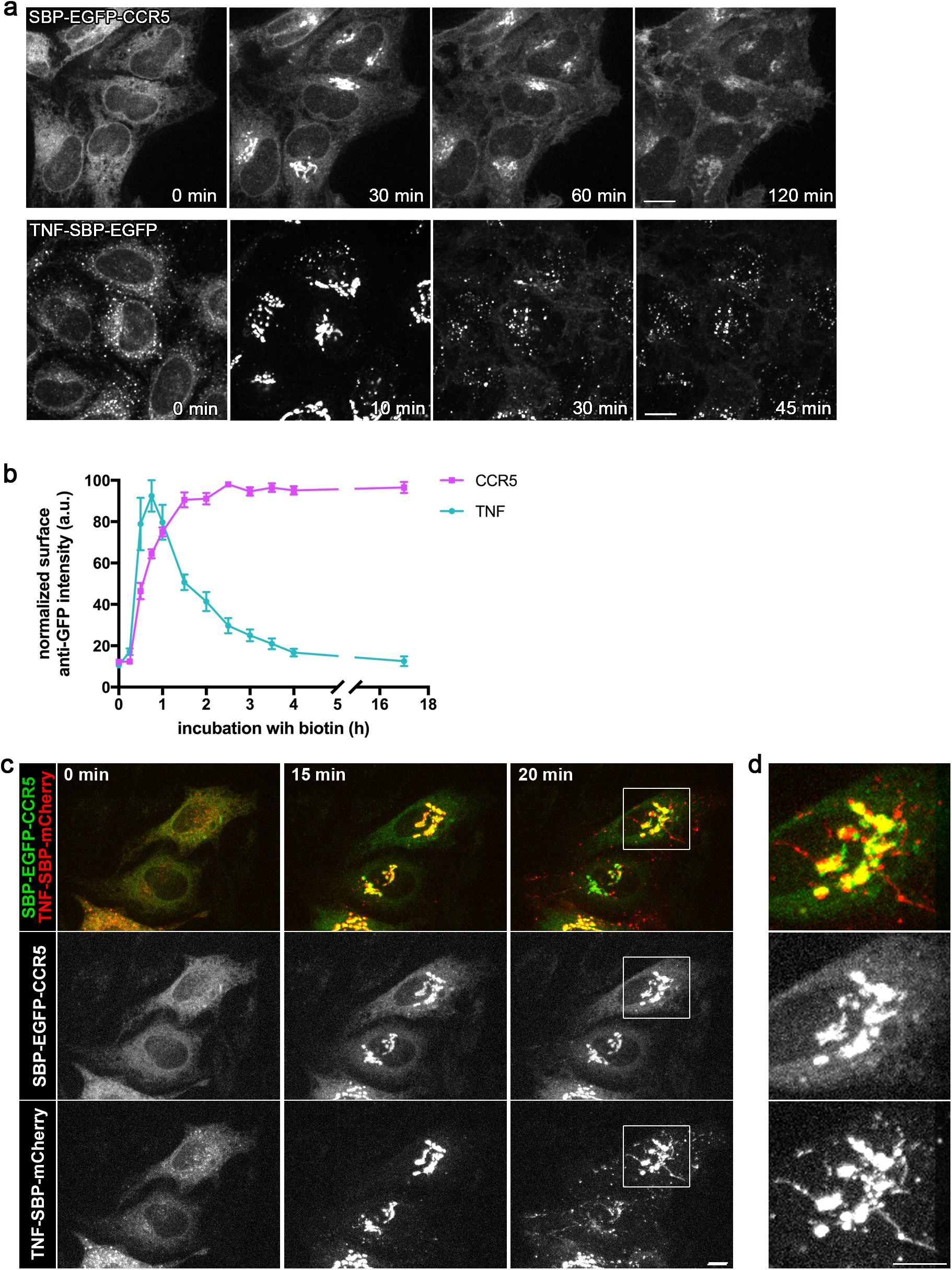
Differential anterograde transport of CCR5 and TNF. **(a)** Synchronized transport of CCR5 (top) and TNF (bottom) in HeLa cells stably expressing Str-KDEL_SBP-EGFP-CCR5 or Str-KDEL_TNF-SBP-EGFP. Trafficking was induced by addition of biotin at 0 min. Scale bar: 10 µm. **(b)** Kinetics of arrival of CCR5 (magenta) or TNF (cyan) to the cell surface after release from the endoplasmic reticulum measured by flow cytometry. Ratio of cell surface signal divided by GFP intensity was used for normalization. The mean ± sem of 3 experiments is shown. **(c)** Dual-color imaging of the synchronized transport of SBP-EGFP-CCR5 and TNF-SBP-mCherry transiently co-expressed in HeLa cells. Streptavidin-KDEL was used as an ER hook. Release from the ER was induced by addition of biotin at 0 min. Scale bar: 10 µm. **(d)** 2.8x magnification of the Golgi complex region is displayed. Scale bar: 10 µm See also Supplementary videos 1-3

### Identification of molecules inhibiting CCR5 secretion by high-content screening

To identify molecules that specifically inhibit the secretion of CCR5, high-content screenings of chemical libraries were conducted. HeLa cells stably expressing either RUSH-adapted CCR5 (Str-KDEL_SBP-EGFP-CCR5) or TNF (Str-KDEL_TNF-SBP-EGFP) were plated in 384-well plates and treated with small molecules from two chemical libraries: a 1,200 FDA-approved drug collection from the Prestwick company and a 2,824 drug collection obtained from the American National Cancer Institute (NCI). Cells were incubated with the molecules at 10 µM for 1h30. As the molecules were dissolved in DMSO, an identical concentration of DMSO was used as negative control. Brefeldin A (BFA), which blocks secretion, and nocodazole, which disrupts microtubules and perturbs Golgi organization, were used as additional controls. In addition, biotin was omitted in some wells to validate the screening procedure and the analysis. Transport to the cell surface was induced by incubation with biotin for 2h (for CCR5) or 45 min (for TNF) according to the previously determined secretion kinetics. The global localization of the cargos was determined using GFP fluorescence while the fraction of the cargo present at the cell surface was quantified by immunolabelling using an anti-GFP antibody on fixed, but non-permeabilized, cells. Nuclei were counterstained with DAPI for imaging and segmentation purposes (**Fig. 2a**). As expected, in the absence of biotin (DMSO without biotin), the GFP signal was visible in the ER for CCR5 and TNF, and almost no surface anti-GFP signal was visible. After incubation with biotin (DMSO with biotin), CCR5 and TNF reached the plasma membrane as shown by cell surface staining (see **Fig. 2d**). Using features obtained from image segmentation, a bioinformatics analysis was conducted to identify molecules from the Prestwick (**Fig. 2b**) and NCI libraries (**Fig. 2c**) that alter different parameters such as, CCR5 or TNF localization and secretion, cell organization, or that induce cell death. Principal component analysis (PCA) and hierarchical clustering, allowed the grouping of conditions where altered secretion was expected (*i.e.* DMSO without biotin, BFA with biotin, and nocodazole with biotin) and to separate them from normal conditions (*i.e.* DMSO with biotin). The same approach was used to identify molecules from the libraries that altered secretion of CCR5 or TNF. Several small molecules prevented secretion of both CCR5 and TNF (‘CCR5 and TNF hit’). In addition to these generic inhibitors, several molecules specifically perturbed the secretion of either CCR5 (‘CCR5 specific hit’) or TNF (‘TNF specific hit’) (**Fig. 2e**). The CCR5 specific hits were then further analyzed.

**Figure 2:**
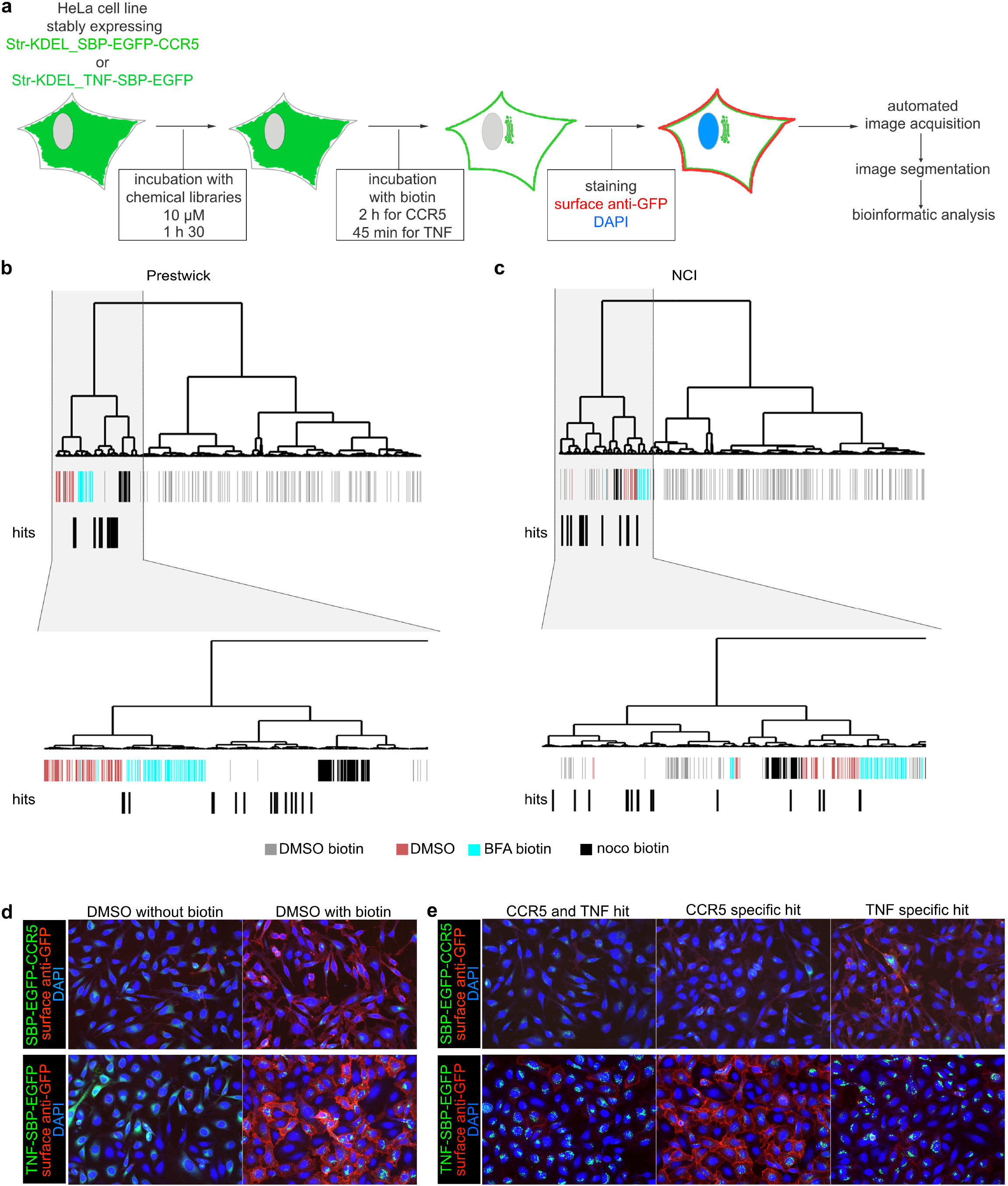
Identification of molecules inhibiting CCR5 secretion by high-content differential screening **(a)** Outline of the chemical screening strategy. **(b)** Clustering of molecules obtained after bioinformatics analysis of the Prestwick chemical library screening of CCR5 secretion. **(c)** Clustering of molecules obtained after bioinformatics analysis of the NCI library screening of CCR5 secretion. Below the dendrogram, each bar corresponds to a well of the plate containing either control molecule or a molecule from the libraries. Molecules identified as hits were shifted one lane below for better visualization. Enlargement of the region containing hits is displayed for better visualization. BFA, brefeldin A; noco, nocodazole. **(d)** Micrographs from screening plates showing controls. DMSO without biotin corresponds to the condition where no secretion occurred and DMSO with biotin corresponds to normal secretion. HeLa cells stably expressing Str-KDEL_SBPEGFP-CCR5 are displayed in the top panels and HeLa cells stably expressing Str-KDEL_TNF-SBP-EGFP are displayed in the bottom panels. Presence of the cargo at the cell surface was detected using labeling with anti-GFP antibodies on nonpermeabilized cells (red). **(e)** Micrographs from screening plates showing the three classes of hits detected. ‘CCR5 and TNF hit’ corresponds to a molecule affecting secretion of both CCR5 and TNF. ‘CCR5 specific hit’ or ‘TNF specific hit’ corresponds to a molecule inhibiting only CCR5 secretion or only TNF secretion, respectively. HeLa cells stably expressing Str-KDEL_SBP-EGFP-CCR5 are displayed in the top panels and HeLa cells stably expressing Str-KDEL_TNF-SBP-EGFP are displayed in the bottom panels. Presence of the cargo at the cell surface was detected using labeling with anti-GFP antibodies on non-permeabilized cells (red).

### Hit validation and specificity over two other chemokine receptors

The 15 strongest CCR5 hits (**Table 1**) were selected for further analysis. As a secondary screen, the effects of the molecules on the trafficking of two other chemokine receptors, CCR1 and CXCR4, were evaluated. CCR5, CCR1 and CXCR4 all belong to the class A subfamily (rhodopsin-like) of GPCR. CCR5 shares several ligands with CCR1 (namely CCL3, CCL4 and CCL5) demonstrating that they are closely related. CXCR4 is a co-receptor that mediates HIV-1 entry into target cells of strains that emerge at an advanced stage of infection. CCR1 and CXCR4 secretion was synchronized using the RUSH assay. The export of CCR1 and CXCR4 secretion were slightly faster than those of CCR5 but, like CCR5, they then remained stably expressed at the cell surface over several hours (**Fig. 3a**). An end-point assay was conducted to measure the effects of the 15 molecules on the secretion of the chemokine receptors (CCR5, CCR1 and CXCR4). The molecules were ranked according to their relative impact on CCR5 secretion. Molecules 1 to 10 reduced the transport of CCR5 to the cell surface by less than 50% while a stronger effect was observed for molecules 11 to 15. In particular, molecules 13, 14, and 15 reduced the cell surface expression of CCR5 by more than 75% (**Fig. 3b**). These three molecules had only moderate effects on the secretion of CCR1 and CXCR4, demonstrating their specificity of action on the transport of CCR5 towards the cell surface (**Fig. 3c, 3d**). Dual color-imaging of the synchronized secretion of CCR5 and CCR1 in a same cell was carried out in both non-treated cells and in cells pre-treated with molecule 13 (Fig. 3e, f). In non-treated cells, CCR5 and CCR1 both exited from the ER and reached the Golgi complex and the cell surface after addition of biotin (**Fig. 3e**). In contrast, when cells were pre-treated with molecule 13, CCR1 reached the cell surface as in non-treated cells, while CCR5 transport was delayed (**Fig. 3f**). CCR5 was still observed in the ER and in the Golgi complex after more than 2h, indicating that molecule 13 inhibits the trafficking of CCR5 at early stage. These experiments demonstrate that molecule 13 is not a broad inhibitor of chemokine receptor transport but rather specifically targets CCR5 transport.

**Table 1:**
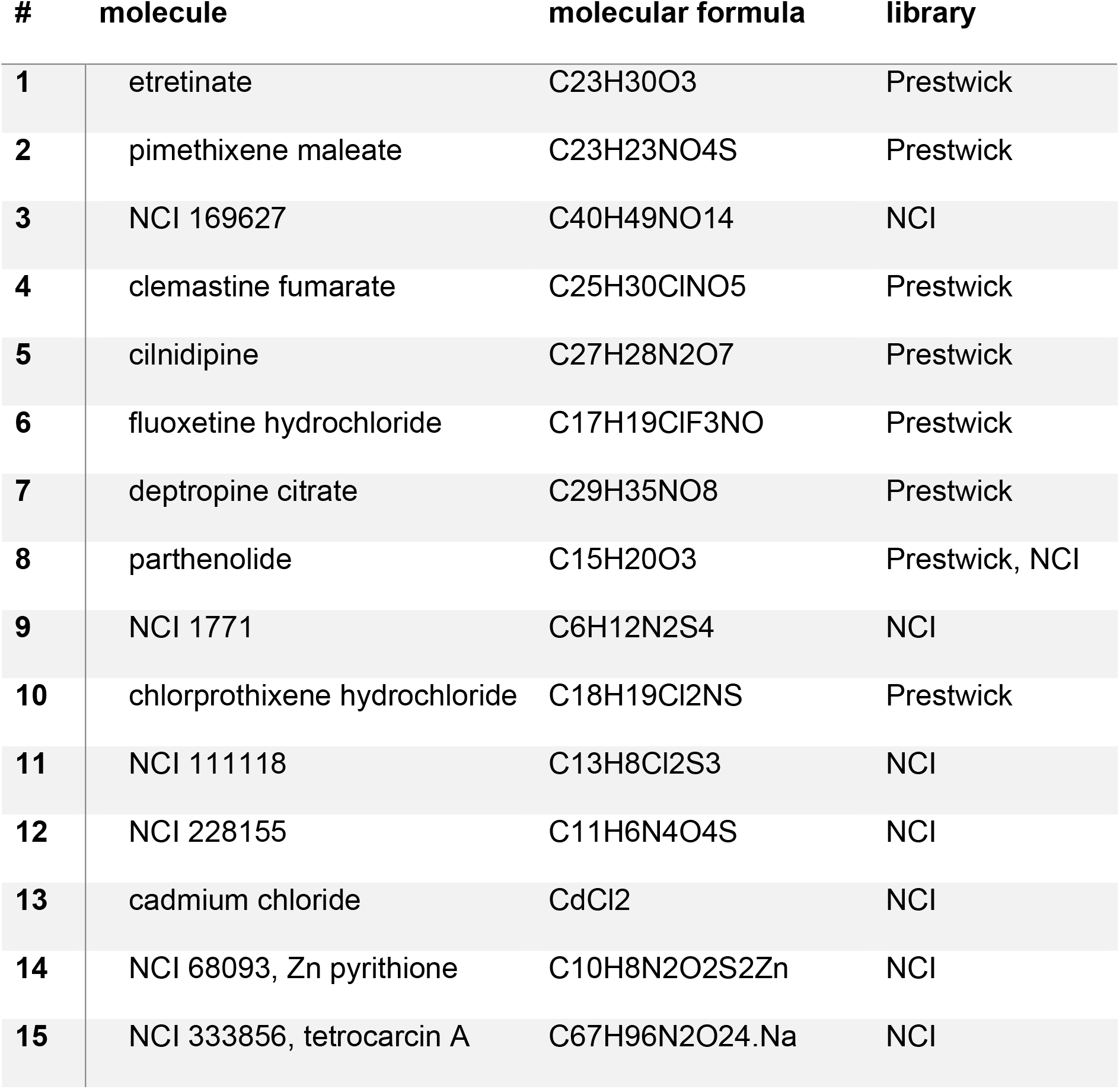
Names and molecular formulas of the molecules.

**Figure 3:**
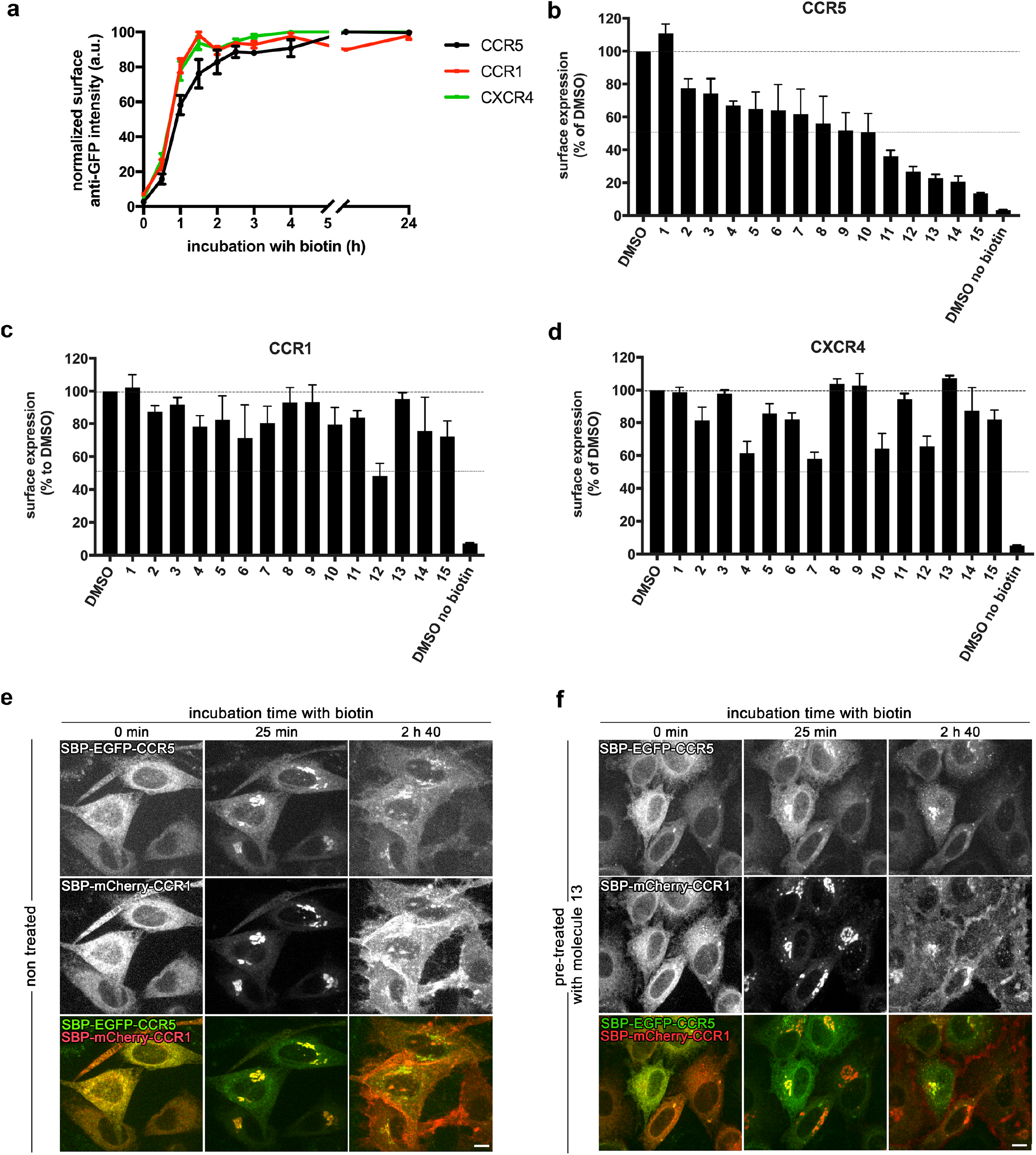
Three small molecules inhibit the secretion of CCR5 and do not affect secretion of CCR1 nor CXCR4 **(a)** Kinetics of synchronized transport to the cell surface of three chemokine receptors, CCR5 (black), CCR1 (red), and CXCR4 (green) using the RUSH assay. Trafficking was induced by addition of biotin at time 0. Surface expression of CCR5, CCR1 and CXCR4 was measured by flow cytometry using an anti-GFP antibody. Ratio of cell surface signal divided by GFP intensity was used for normalization. The mean ± sem of 3 experiments is shown. End-point measurement (2h) of the effects of the hit molecules identified in the primary screen on the trafficking of CCR5 (for validation) **(b)**, of CCR1 **(c)** and of CXCR4 **(d)** in HeLa cells. Cells were pre-treated for 90 min with molecules at 10 µM. The amount of cargo present at the cell surface 2h after release from the ER was measured by flow cytometry using an anti-GFP antibody. Ratio of cell surface signal divided by GFP intensity was used for normalization. The mean ± sem of 3 experiments is shown. Real-time synchronized secretion of CCR5 and CCR1 was monitored using dual-color imaging in both non-treated HeLa cells **(e)** and in HeLa cells pre-treated for 90 min with 10 µM of molecule 13 **(f)**. Scale bar: 10 µm.

### Molecules 13 and 14 inhibit the secretion of CCR5 via its cysteine-containing cytoplasmic tail and its palmitoylation

Little is known about the key players controlling the secretion of CCR5 and no specific pathway has been reported so far. CCR5 is a seven transmembrane domain protein with its amino-terminal extremity facing the luminal/extracellular space and its carboxy-terminal extremity in the cytoplasm (**Fig. 4a**). As the carboxy-terminal domains of CCR1 and CCR5 differ, the involvement of the cytoplasmic tail of CCR5 in mediating the effects of molecules 13, 14 and 15 on receptor secretion was examined. Chimeric receptors were constructed by exchanging the cytoplasmic tails of CCR5 and CCR1. (**Fig. 4b**). Using the RUSH assay, it was determined that the CCR5-CCR1tail and CCR1-CCR5tail were able to reach the cell surface with kinetics similar to CCR5 and CCR1 (**Fig. 4c**). Incubation of cells expressing either these chimeras or the controls, revealed that molecules 13, 14 and 15 affected the secretion of the chimeras in different ways. Molecules 13 and 14 inhibited the secretion of the constructs bearing the cytoplasmic tail of CCR5 (*i.e.* CCR5wt and CCR1-CCR5tail) by more than 40 %. They were however, quite inefficient against receptors bearing CCR1 tail (**Fig. 4d**). In contrast, the transport of both chimeras (CCR5-CCR1tail and CCR1-CCR5tail) was inhibited after exposure to molecule 15 by more than 40 %.

**Figure 4:**
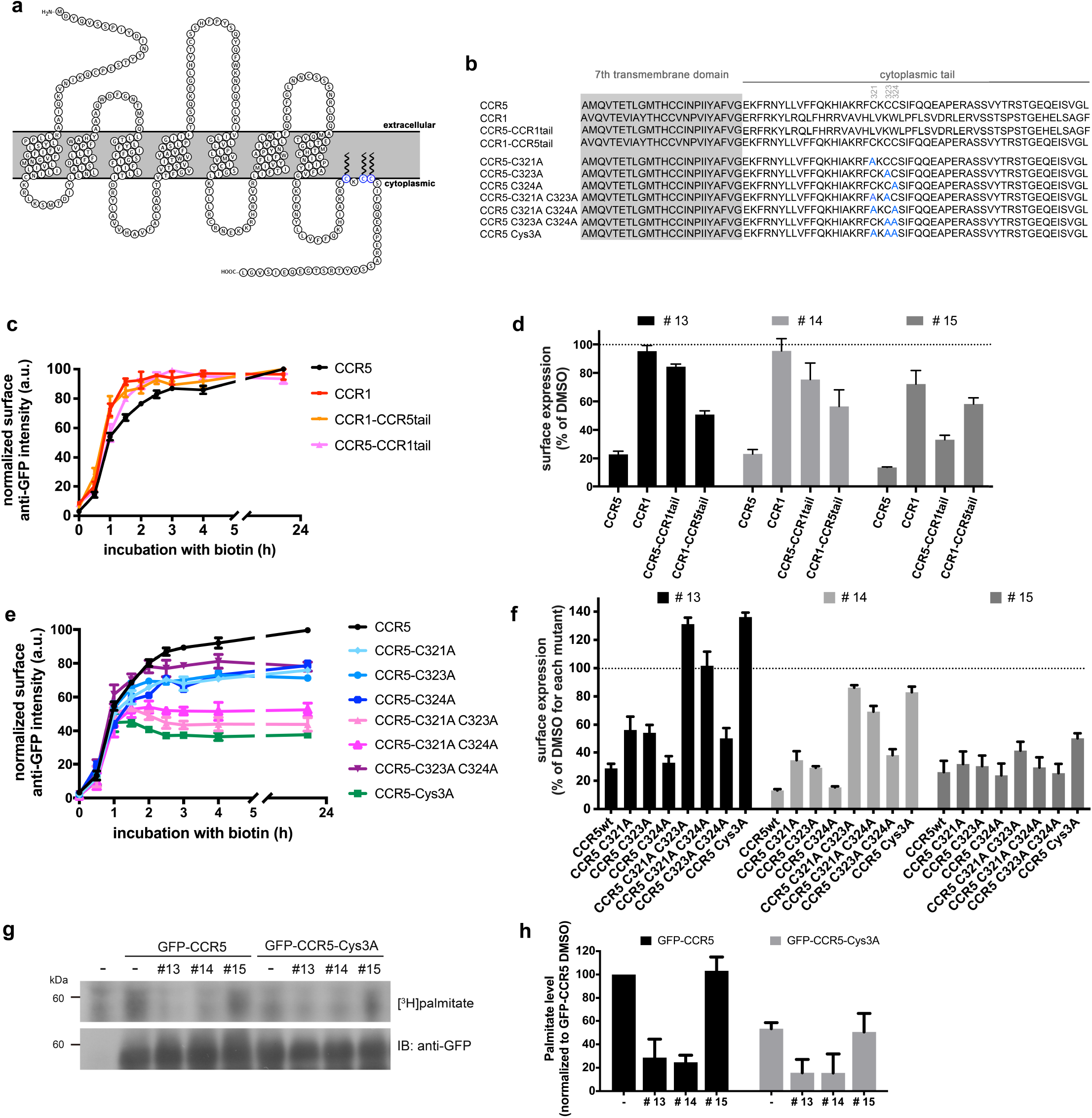
Molecules 13 and 14 inhibit CCR5 secretion via its cytoplasmic tail and inhibit palmitoylation. **(a)** Schematic representation of CCR5 with three palmitoylated cysteine residues indicated in blue. **(b)** Amino-acid sequence of CCR5, CCR1, their chimeras, and the cysteine to alanine mutants used in this study. **(c)** Kinetics of the synchronized transport of CCR5/CCR1 chimeras to the cell surface measured by flow cytometry in HeLa cells transiently expressing SBP-EGFPCCR5 (black), SBP-EGFP-CCR1 (red), SBP-EGFP-CCR1-CCR5tail (orange) and SBP-EGFP-CCR5-CCR1tail (magenta). Release from the ER was induced by addition of biotin at time 0. The mean ± sem of 3 experiments is shown. **(d)** End-point measurement (2h) of the effects of molecules 13, 14, and 15 on the secretion of the CCR5/CCR1 chimeras. HeLa cells transiently expressing either, CCR5, CCR1, CCR5-CCR1tail, or CCR1-CCR5tail were pre-treated for 1.5 h with molecules at 10 µM. The amount of cargo present at the cell surface 2h after release from the ER was measured by flow cytometry using an anti-GFP antibody. The mean ± sem of 3 experiments is shown. **(e)** Kinetics of the synchronized transport of CCR5 cysteine mutants to the cell surface measured by flow cytometry in HeLa cells transiently expressing the corresponding SBP-EGFP-CCR5 wt or mutant construct. Release from the ER was induced by addition of biotin at time 0. The mean ± sem of 3 experiments is shown. **(f)** End-point measurement (2h) of the effects of molecules 13, 14 and 15 on the secretion of CCR5 cysteine mutants. HeLa cells transiently expressing the cysteine mutants were pre-treated for 1.5 h with molecules at 10 µM. The amount of cargo present at the cell surface 2h after release from the ER was measured by flow cytometry using an anti-GFP antibody. The mean ± sem of 3 experiments is shown. (g,h) Quantification of palmitoylation of either GFP-CCR5 or GFP-CCR5 Cys3A transiently expressed in HEK293T cells. [3H] palmitate was incorporated for 4h in presence of the compounds following a pre-treatment with DMSO (-), molecules 13, 14 and 15 for 30 min. A representative autoradiogram and immunoblot are shown (g) and the mean ± sem of 4 experiments is shown. See also Figure S1

The comparison of the results obtained with chimeras, CCR1-CCR5tail and CCR5-CCR1tail, indicated that the presence of the cytoplasmic tail of CCR5 was necessary for the reduction of trafficking induced by molecules 13 and 14. The cytoplasmic tail of CCR5 contains three cysteine residues that require palmitoylation to ensure efficient secretion ^18, 19^. In contrast, the CCR1 cytoplasmic tail does not contain palmitoylated cysteine residues. To further study the role of the three cysteine residues on the cytoplasmic tail of CCR5 in mediating the effect of molecules 13 and 14, a series of mutants were created with cysteine substituted by alanine, independently or in a combinatorial way (**Fig. 4b**). The kinetics of secretion of the CCR5 cysteine mutants were consistent with previous reports ^18, 19^. The secretion of the single cysteine mutants was decreased by 20%, while the secretion of the double and triple cysteine mutants was decreased by 50%, with the exception of the CCR5 C323A-C324A mutant that was decreased by only 20% (**Fig. 4e**). We then tested the effect of the small molecules on the transport of the cysteine mutants. Molecule 13 did not reduce the secretion of CCR5 C321A-C323A, CCR5 C321AC324A and CCR5 Cys3A, while other mutants were still affected (e.g. CCR5 C324A) (**Fig. 4f**). Molecule 14, while more potent than molecule 13, displayed the same relative dependence on the different mutations. In contrast, the inhibitory effect of molecule 15 was not affected by cysteine mutation and transport to the cell surface decreased by at least 50 % for all mutants.

Because these cysteine residues were reported to be targets of palmitoylation, we directly assessed the effects of molecules 13, 14 and 15 on CCR5 palmitoylation in cells. In vivo metabolic labeling with radioactive palmitate of cells expressing either GFP-CCR5 or GFP-CCR5 Cys3A was performed in control cells or in cells treated by the molecules (Fig. 4g, 4h). Molecules 13 and 14 decreased the level of palmitoylated GFP-CCR5 by 70%. In agreement with our previous results suggesting that molecule 15 does not target CCR5 tail, it did not affect the palmitoylation level of GFP-CCR5 compared to DMSO control. As expected, GFPCCR5 Cys3A showed a reduced level of palmitoylation (about 50%) compared to GFP-CCR5. However, radioactive signal was still detected in cells expressing GFPCCR5 Cys3A, suggesting residual palmitoylation on cysteines other than Cys 321, 323 and 324. Interestingly, this residual signal was also decreased after incubation with molecules 13 and 14, while molecule 15 had no effect.

We looked for pamitoyltransferase able to modify CCR5 and showed. DHHC3, DHHC7 and DHHC15 were found to induce palmitoylation of CCR5 (Fig. S1a). Autopalmitoylation of DHHC3 and DHHC7 was inhibited to about 50 % following incubation with molecules 13 and 14, whereas molecule 15 had no effect (Fig. S1b, S1c). These results suggest that molecules 13 and 14 may inhibit autopalmitoylation of DHHCs responsible for CCR5 palmitoylation and consequently palmitate transfer to CCR5.

Altogether, our results indicate that molecules 13 and 14 may share a similar mode of action inducing a strong reduction of CCR5 palmitoylation. In contrast, molecule 15 seems to affects the secretion of CCR5 by another, still elusive, mechanism.

### Inhibition of HIV-1 infection in primary human macrophages

The effects of molecules 13, 14 and 15 on the cell surface expression of CCR5 in human monocyte-derived macrophages (hMDM) after overnight treatment were studied. These molecules, alone or in combination, induced a small but significant decrease of CCR5 cell surface expression (19.1 to 23.8 %) compared with DMSO treatment. The cell surface expression of CXCR4 however, was not significantly modified under the same conditions (p>0.05; **Fig. 5a, 5b**). The treatment with the three molecules, alone or in combination, did not induce major cytotoxicity as compared with DMSO (**Fig. 5c**). Therefore, as observed in HeLa cells, molecules 13, 14, and 15 selectively impacted CCR5 in primary macrophages.

**Figure 5:**
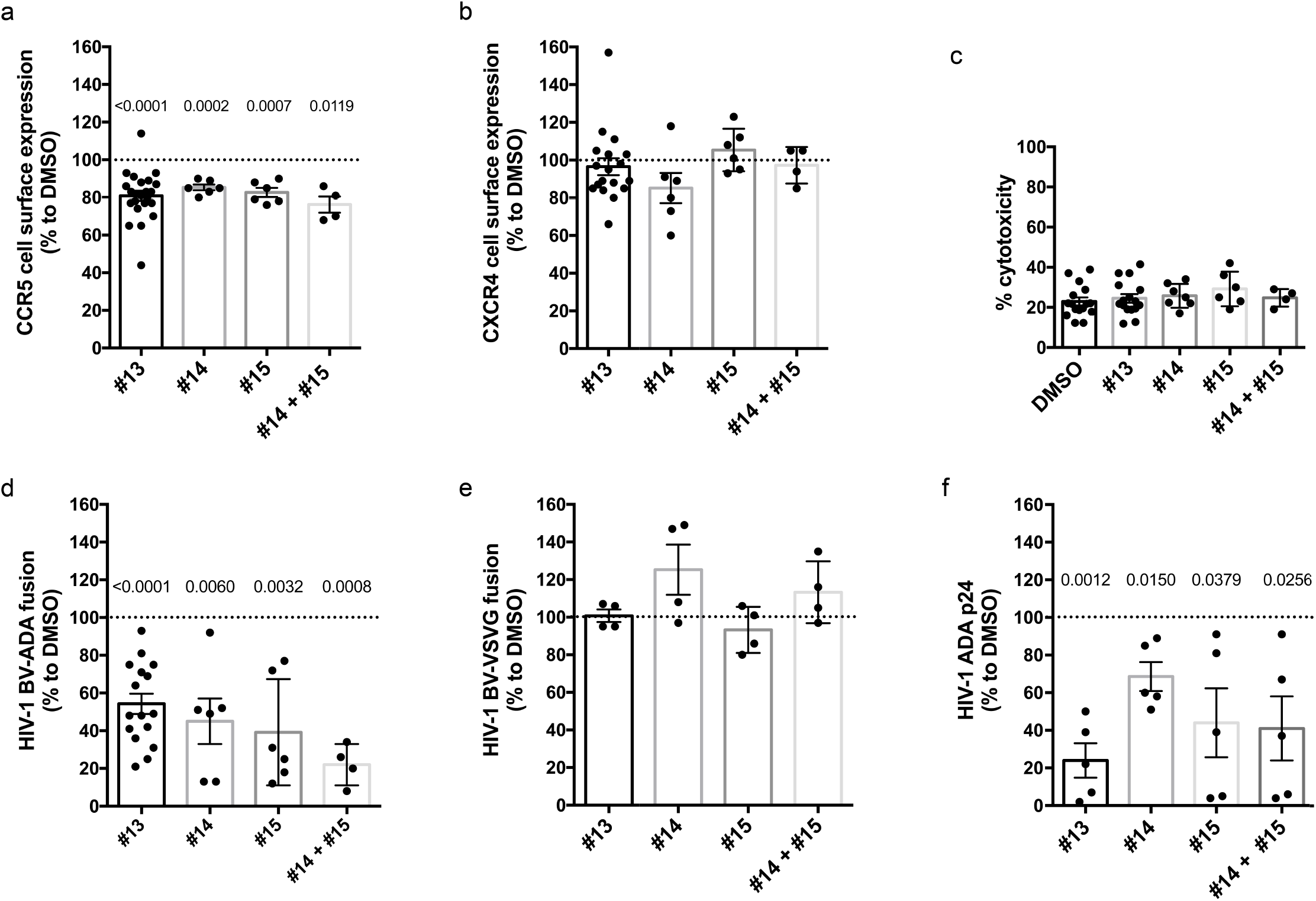
Treatment with molecules 13, 14 and 15 decreases HIV-1 R5 infection in human macrophages. Primary human macrophages differentiated for 4 days with rhM-CSF were treated during 18 h with molecule 13 at 10 µM; molecule 14 at 3 µM; molecule 15 at 1 µM and molecules 14 and 15 at 1 µM (or DMSO at 0.1%). Cell surface expression of CCR5 (**a**) or CXCR4 (**b**) was measured by flow cytometry with specific antibodies. (**c**) Cytotoxicity was evaluated by measuring LDH release in human primary macrophages after 18 h pre-treatment with molecules or DMSO as a control. Inhibition of fusion of HIV-1_ADA_ (**d**) or HIV-1 _VSVG_ (**e**) containing BlaM-vpr (BV) with CCF2/AM primary macrophages mediated by compounds. Total p24 amount (supernatants + lysates) of HIV-1_ADA_-infected macrophages measured by ELISA (**f**). Each black point represents one donor analyzed independently. Statistical analyses were performed using PRISM software. A one sample T-test was applied and significant p-values (< 0.05) are indicated for each treatment compared to DMSO in (**a**), (**d**) and (**f**). The absence of a p-value indicates that the results were not significantly different. Error bars correspond to SEM.

The capacity of HIV particles to fuse with cells after overnight treatment with the same molecules was then investigated using the BlaM-Vpr fusion assay ^20^. The fusion of HIV-1 ADA particles with hMDM treated with molecules 13, 14 and 15 was strongly decreased (by 45.7 to 78.0 %) relative to those treated with DMSO (**Fig. 5d**). Under the same conditions, the entry of a VSV-G pseudotyped virus, used as a CCR5-independent control, was not affected (**Fig. 5e**). Treatment with both molecules 14 and 15 led to further perturbation of viral entry. These data indicate that a small reduction of neo-synthesized CCR5 expressed at the cell surface is sufficient to significantly impact R5-tropic HIV-1 entry into human macrophages. To further assess whether the perturbation of viral entry was sufficient to alter the viral cycle and production, the total amount of p24 capsid protein produced by macrophages was quantified. Viral production and secretion were both strongly reduced, by 31.4 to 76.0 %, in particular in cells treated with molecule 13. (**Fig. 5f**). Taken together, these results demonstrate that molecules specifically reducing CCR5 secretion at the cell surface impair HIV-1 infection of human macrophages.

## Discussion

The development of many pathologies rely on the efficient intracellular transport of proteins. Transport to the cell surface is particularly important as adhesion proteins, channels, protease or receptors for example have to reach the plasma membrane to fulfill their functions. We thus reasoned that, instead of looking for molecules able to perturb their function, we may target their transport pathway to prevent their normal expression at the cell surface, hence exploring a novel therapeutic option. This option has been under-exploited for at least two reasons:

On the one hand, it has long been thought that the diversity of secretory routes was low and that the bulk flow of membranes was responsible for most non-specialized pathways. We now know that the diversity of pathways is high with several coats, adaptors, Golgi matrix proteins or molecular motors active at the same transport stage. In addition, not only the molecular machinery of transport may be targeted, the cargo itself may be targeted. Perturbation of its folding, modifications, interaction with transport partners or membrane partitioning may perturb, or prevent, its transport.

On the other hand, as compared to the power and precision of the study of endocytosis and retrograde pathways, the diversity of the secretory pathway has long been hard to study and target. Quantitative monitoring of the transport of proteins was only possible for selected proteins and assays were hardly amenable to screening. The development of the RUSH assay ^16^ that allows to cope with the diversity now allows us to overcome this limitation and specifically screen for inhibitory molecules.

To validate our hypothesis; we used the RUSH assay to screen for small molecules able to inhibit specifically the transport of CCR5 to the cell surface. CCR5 is indeed essential for R5-tropic HIV-1 strain infections of human cells and represents a valuable therapeutic target. Individuals devoid of CCR5 expressed at the cell surface are resistant to HIV-1 infection as CCR5 while people heterozygous for this deletion show a reduction of CCR5 cell surface expression and slower progression of HIV infection ^21^. Importantly, the absence of functional CCR5 seems not to be deleterious to these individuals, though increased susceptibility to infections, such as West Nile virus ^22, 23^ and tick-borne encephalitis, ^24^ have been reported.

Several large-scale screens were conducted to identify protein regulators of HIV-1 infection, but none of them led to the identification of CCR5 secretion regulators ^25-28^. Little is known about the molecular players that regulate the secretion of CCR5. The involvement of the small GTPases, Rab1, Rab8 and Rab11 has been proposed ^29^. Rab43 was recently reported to play a role in the export of several class A GPCRs from the ER ^30^, though its role in controlling the transport of CCR5 was not evaluated. The importance of palmitoylation of CCR5 was also revealed ^18, 19^. However, the palmitoyltransferase responsible for palmitoylation of CCR5 remains unknown.

RUSH-based differential screening, using TNF as a control reporter, allowed to identify a set of inhibitory molecules. In particular, three molecules that strongly perturbed CCR5 transport were found to have no, or only moderate, effects on the secretion of the closely related CCR1 and CXCR4. Molecules 13, 14 and 15 were active on endogenous CCR5 secretion and reduced HIV-1 infection of human macrophages isolated from donors. Molecules 13, 14 and 15 are cadmium chloride (CdCl_2_), zinc pyrithione (C_10_H_8_N_2_O_2_S_2_Zn) and tetrocarcinA (C_67_H_96_N_2_O_24_), respectively. Cd and zinc pyrithione were found to share similar mechanisms of action since they both rely on the C-terminal cytoplasmic tail of CCR5 and in particular the cysteine residues in position 321, 323 and 324 and they both inhibit CCR5 palmitoylation. At the molecular level, Cd affects protein function by binding to thiol groups. Direct binding of Cd to cysteine residues may prevent efficient palmitoylation of CCR5, hence affecting its transport, as previously reported ^18, 19^. Alternatively, palmitoyltransferase may represent the target of these molecules. Palmitoylation occurs in a two step mechanism. First, palmitoyl-CoA is transferred to the DHHC cysteine-rich domain leading to autoacylation of the palmityoyltransferase ^31^. DHHC contains bound Zn ^32^. In the second step, the palmitoyl group is transferred to the cysteine residues of the target protein. Cd is known to displace essential metals like Zn in metalloproteins (see ^33^ for review). Zn pyrithione is an antifungal and antibacterial zinc chelator. As these two molecules may affect Zn-dependent enzymes, it is tempting to propose that Zn pyrithione and Cd bind to, and perturb, the palmitoyltransferase responsible for palmitoylation of CCR5.

Tetrocarcin A represent another class of CCR5 inhibitory molecule because it does not depend on the C-terminal tail and does not impact palmitoylation. It is an antibiotic ^34^ which antagonizes Bcl-2 anti-apoptotic function ^35^. The putative mode of action of tetrocarcin A on the trafficking of CCR5 remains unclear. It has been reported that Tetrocarcin A induces ER stress in B-chronic lymphocytic leukemia cells ^36^. However, cell treatment with tetrocarcin A did not lead to a general ER stress that would affect the secretory pathway. While tetrocarcin A inhibited the secretion of CCR5, it did not affect TNF, CCR1 or CXCR4 transport. Tetrocarcin A may affect the conformation of CCR5 by an as yet unknown mechanism or perturb interaction with the proteins that regulate CCR5 transport. Further studies are required to obtain a clearer view of tetrocarcin A’s mechanism of action but it may represent an interesting novel class of CCR5 inhibitory molecule.

The three molecules identified in our study exhibited inhibitory effects on HIV-1 infection for R5-tropic viruses both at the level of virus entry and viral particle production in human macrophages. Of major concern in anti-HIV therapies, particularly those targeting a receptor such as CCR5, is the emergence of escape viruses. To date, Maraviroc is the only approved anti-HIV therapy targeting CCR5. Reports of the emergence of resistant viruses have since been published. The outgrowth of CXCR4-tropic viruses was observed in some patients ^37^. For this reason, only patients with R5 tropic HIV-1 strains are eligible for treatment with Maraviroc. In other patients, viral glycoproteins accumulated mutations enabling interaction with CCR5 bound to Maraviroc ^38, 39^. Targeting the host secretory pathway may avoid the emergence of such escape viruses. The molecules identified in this study induced a decrease in secretion of CCR5 resulting in a decrease in expression at the cell surface. If mutations were to arise in R5-tropic viruses, they would not be able to induce normal secretion of CCR5 and restore infection. This is a clear advantage over treatments based on competition or allosteric modifications. Combinatorial treatment may also reduce the amount of CCR5 at the cell surface and may therefore improve the efficacy of blocking antibodies or of Maraviroc.

In conclusion, three molecules that impact CCR5 secretion and reduce R5-HIV infection of human macrophages were identified using chemical screening. Two inhibitory molecules highlighted the importance of the C-terminal tail of CCR5 for its transport and its value to develop anti-HIV strategies, while a third one may uncover a novel class of inhibitory molecules. More generally, our study confirms our model that proposes that the diversity of secretory routes can be exploited to identify molecules that specifically affect the transport of a given receptor. Targeting the transport and nor the function of target protein may thus represent a novel therapeutic paradigm.

## Methods

### Cells

HeLa cells were cultured in Dubelcco’s modified Eagle medium (DMEM) (Thermo Fisher Scientific) supplemented with 10% Fetal Calf Serum (FCS, GE Healthcare), 1 mM sodium pyruvate and 100 µg/ml penicillin and streptomycin (Thermo Fisher Scientific).

HeLa cells stably expressing Str-KDEL as a hook and either SBP-EGFP-CCR5 or TNF-SBP-EGFP as a reporter were obtained by transduction with lentiviral particles produced in HEK293T. A clonal population was then selected using puromycin resistance and limiting dilution.

Human primary macrophages were isolated from the blood of healthy donors (Etablissement Français du Sang Ile-de-France, Site Trinité, #15/EFS/012) by density gradient sedimentation in Ficoll (GE Healthcare), followed by negative selection on magnetic beads (Stem cells, Cat n°19059) and adhesion on plastic at 37°C for 2 h. Cells were then cultured in the presence of complete culture medium [RPMI 1640 supplemented with 10 % Fetal Calf Serum (FCS) (Eurobio), 100 µg/ml streptomycin/penicillin and 2 mM L-glutamine (Invitrogen/Gibco)] containing 10 ng/ml rhM-CSF (R&D systems) ^40^ for 4 - 5 days.

### Plasmids and transfection

The DNA sequences corresponding to human CCR5 (UniProt P51681), CCR1 (Uniprot P322246) and CXCR4 (Uniprot P61073) were purchased either as synthetic genes (Geneart, Thermo) or as cDNA (Openbiosystems). They were cloned into RUSH plasmids downstream of Str-KDEL_IL2ss-SBP-EGFP or Str-KDEL_IL2ss-SBP-mCherry using FseI and PacI restriction enzymes ^16^. Str-KDEL_TNF-SBPEGFP and Str-KDEL_TNF-SBP-mCherry plasmids have been described elsewhere ^16^. The CCR5-CCR1tail and CCR1-CCR5tail chimeras were generated from synthetic genes (Geneart, Thermo) and cloned between FseI and PacI restriction sites. Mutations from cysteine to alanine in CCR5 tail were generated either by PCR assembly or by the insertion of small, synthetic DNA fragments (gBlock from IDT DNA). Protein sequences of chimeras and mutants are depicted in Fig. 4b. The GFPCCR5 and GFP-CCR5-Cys3A bear the IL-2 signal peptide upstream of GFP and either CCR5 wild-type or CCR5-Cys3A downstream of GFP. A modified version of pEGFP (Clontech) was used for their generation. All plasmids used in this study were verified by sequencing.

HeLa cells were transfected using calcium phosphate as described previously ^41^.

The HIV-1_ADA_ provirus plasmid (pHIV-1_ADA_) expressing the *env* gene of the HIV-1 R5-tropic strain ADA has been described elsewhere ^42^. pNL4.3D pNL4.3f was a gift from P. Benaroch (Institut Curie, Paris, France). The plasmid expressing the *env* gene of VSVG (pEnv_VSVG_) was a gift from S. Benichou (Institut Cochin, Paris, France).

### High-content automated chemical screening

Chemical compounds were purchased from Prestwick Chemicals (Illkirch, France) corresponding to 1,200 approved drugs (FDA, EMA and other agencies) dissolved in dimethyl sulfoxide (DMSO) at 10 mM. A second library of 2,824 compounds was kindly provided by the National Cancer Institute - NCI chemical libraries as follows: Diversity Set III, 1,596 compounds; Mechanistic Set, 879 compounds; Approved Oncology Drugs set II, 114 agents and Natural products set II, 235 agents. All NCI stock compounds were received in DMSO at a concentration of 10 mM except for Mechanistic Set (at 1 mM) (in a 96-well plate format). All libraries were reformatted in-house in 384-well plates. Brefeldin A (BFA) and nocodazole were purchased from Sigma and used as control molecules.

For compound screening, cells (5.0 × 10^3^ per well) were seeded on black clear bottom 384-well plates (ViewPlate-384 Black Perkin Elmer) in 40 µL of complete medium. The screen was performed at the similar early cell passages (±2) for both replicates. Twenty-four hours after cell seeding, compounds were transferred robotically to plates containing cells using the TeMO (MCA 384) (TECAN) to a final concentration of 10 µM and 0.5% of DMSO. Controls were added to columns 1, 2, 23 and 24 of each plate. After 90 min of compound incubation, cells were treated with 40 µM biotin for 45 minutes (for TNF) or 120 minutes (CCR5) at 37°C. Compound screens were performed in two independent replicate experiments at the BioPhenics Screening Laboratory (Institut Curie).

Cells were processed immediately after biotin treatment for immunofluorescence. Briefly, cells were fixed with 3% paraformaldehyde for 15 min and quenched with 50 mM NH_4_Cl in phosphate buffered saline (PBS) solution for 10 min. For cell surface labelling, cells were incubated with anti-mouse GFP (1:800, Roche, Cat N° 814 460 001) diluted in 1% BSA blocking solution for 45 min. Cells were then washed with PBS and incubated for 1 h with Cy3-conjugated anti-mouse (1:600, Jackson Immunoresearch, Cat N° 715-165-151). Nuclei were counterstained with DAPI (Life Technologies) for 45 min.

Image acquisition was performed using an INCell 2200 automated high-content screening fluorescence microscope (GE Healthcare) at a 20X magnification (Nikon 20X/0.45). Four randomly selected image fields were acquired per wavelength, well and replicate experiment. Image analysis to identify cells presenting predominantly cell surface or intra-cellular CCR5 and/or TNF localization was performed for each replicate experiment using the Multi Target analysis application module in the INCell analyzer Workstation 3.7 software (GE Healthcare). Results were reported as mean values from four image fields per well.

### Bioinformatics analysis of the screens

Fields with less than 50 cells were filtered out after image segmentation. All cell features extracted from the image analysis step were normalized within each plate by subtracting the median value of control wells containing DMSO and biotin and dividing by the median absolute deviation of the same controls.

Principal Component Analysis (PCA) was applied to normalized data for each dataset separately as a de-noising method. From this PCA, the coordinates of wells in the subspace, defined by principal component with eigen value greater than one, were used to compute the euclidean distance between wells. Hierarchical clustering with Ward’s agglomerative criterion was then applied on these distances. Two groups were identified from the clustering. The group with the positive controls was defined as the hits list.

All analyses were performed using R statistical software.

### Measurement of transport to the cell surface by flow cytometry

1×10^6^ cells expressing RUSH constructs per condition were treated with 40 µM biotin to induce trafficking of the reporter and incubated at 37°C. At the desired time point, cells were washed once with PBS supplemented with 0.5 mM EDTA and incubated with PBS supplemented with 0.5 mM EDTA for 5 min at 37°C. Plates containing cells were then put on ice. Cells were resuspended and transferred to ice-cold tubes for centrifugation at 300 g for 5 min at 4°C. Cell pellets were resuspended in cold PBS supplemented with 1% fetal calf serum for blocking and incubated for at least 10 min. After centrifugation, cell pellets were incubated in a solution of Alexa Fluor 647-coupled anti-GFP (BD Pharmingen, Cat N° 565197) prepared in PBS supplemented with 1% serum for 40 min on ice. Cells were washed 3 times in cold PBS+1% serum and fixed with 2% paraformaldehyde (Electron Microscopy Sciences) for 15 min. Cells were washed twice with PBS prior to acquisition with Accuri C6 flow cytometer. The intensity of the GFP signal (FL1) and of the Alexa Fluor® 647-conjugated antibody (FL4) was measured on GFP-positive cells. The FL4 signal was divided by the FL1 signal to normalize for the transfection level. The FL4/FL1 ratio for each condition was then normalized to the DMSO control.

### Real-time imaging of the synchronized secretion

HeLa cells expressing either stably or transiently RUSH constructs were grown on 25 mm glass coverslips. Prior to imaging (after treatment), coverslips were transferred to an L-shape tubing equipped Chamlide chamber (Live Cell Instrument). Trafficking was induced by exchanging Leibovitz medium (Life Technologies) with pre-warmed Leibovitz medium supplemented with 40 µM of biotin (Sigma). Imaging was performed at 37°C in a thermostat controlled chamber using an Eclipse 80i microscope (Nikon) equipped with a spinning disk confocal head (Perkin) and an Ultra897 iXon camera (Andor). Image acquisition was performed using MetaMorph software (Molecular Devices). Maximum intensity projections of several Z-slices are shown (Fig. 1a,c and Fig. 3e, f).

### Measurement of CCR5 palmitoylation

HEK293T cells were transfected with GFP-CCR5 or GFP-CCR5 Cys3A mutant. 24 h after transfection, cells were labeled with [3 H]palmitate (0.5 mCi/ml) for 4 h. Cells were incubated with 10 µM of individual compounds for 30 min before the labeling and for 4h together with [3 H]palmitate. For compound (-), DMSO was added. For the fluorography, after SDS-PAGE of cell lysates, the gels were exposed for 11-16 days. N = 4 independent experiments. The mean ± sem is shown.

### Antibodies and reagents

The following antibodies were used: Alexa Fluor® 647-coupled anti-GFP (BD Pharmingen, Cat N° 565197), Alexa Fluor® 647 rat IgG2a, κ isotype control (Cat. N° 400526); Alexa Fluor® 647 anti-human CD195 (Cat. N° 313712) from BioLegend; PE mouse IgG1, κ isotype control (Cat. N° 550617); PE mouse IgG2a, κ isotype control (Cat. N°553457); FITC mouse IgG2a, κ isotype control (Cat. N°555573); FITC mouse IgG1 κ isotype control (Cat. N°555748); PE mouse anti-human CD184 (Cat. N°555974); FITC mouse anti-human CD4 (Cat. N°555346); FITC mouse anti-human CD3 (Cat. N°555332); PE mouse anti-human CD11b/Mac1 (Cat. N°555388) from BD Biosciences.

The amount of lactate dehydrogenase (LDH) released into supernatants was quantified using Pierce™ LDH Cytotoxicity assay kit (ThermoFisher Scientific, Cat. N°88953). Alliance® HIV-1 p24 ELISA kit (PerkinElmer®, Cat. N°NEK050) was used to determine the amount of capsid p24 protein in supernatants and cell lysates. LDH and p24 quantification were performed following manufacturer’s instructions.

### Flow cytometry analysis of receptor surface expression in macrophages

Cells were washed once with cold PBS and detached using 2 mM EDTA in PBS on ice. To analyze cell surface expression of receptors, cells were washed with PBS and stained with antibodies (see above) for 1 h on ice in PBS + 2 % Fetal Calf Serum. Cells were then washed twice in cold PBS and analyzed by flow cytometry (Accuri™ C6, BD Bioscience). Propidium iodide (PI) (Sigma-Aldrich, Cat. N°P4864), diluted in cold PBS at 0.1 µg/ml, was added to cells just before analysis.

### Viral production and HIV-1 infection

#### - Viral stocks

HIV-1 particles or pseudo-particles containing BlaM-Vpr were produced by cotransfection of HEK293T cells with proviral plasmids (pHIV-1_ADA_ or pNL4.3ΔEnv + pEnv_VSVG_), pCMV-BlaM-vpr encoding β-lactamase fused to the viral protein Vpr and pAdvantage, as described elsewhere ^20^. After 48 h of culture at 37°C, the virus-containing supernatant was filtered and stored at −80°C. Pseudo-particles containing BlaM-Vpr were ultracentrifuged at 60,000 × g for 90 min at 4°C on a sucrose cushion (20%). The virion-enriched pellet was resuspended in PBS and aliquoted for storage at −80°C. The amount of p24 antigen in the supernatants was quantified using an ELISA kit (PerkinElmer). HIV-1 infectious titers were also determined in HeLa TZM-bl cells (LTRlacZ, NIH reagent program) by scoring beta-lactamase-positive cells 24 h after infection, as described previously ^43^.

#### - BlaM-vpr Viral Fusion assay

After 18 h incubation with compounds, 1.5 x 10^5^ primary macrophages were inoculated with the BlaM-Vpr-containing viruses (15 ng p24 Gag) by 1 h spinoculation at 4°C and incubated 2.5 h at 37°C. Cells were then loaded with CCF2/AM, the BlaM-Vpr substrate (2 h at RT), and fixed. Enzymatic cleavage of CCF2/AM by beta-lactamase ^20^ was measured by flow cytometry (LSR II, BD) and data were analyzed with FACSDiva software. The percentage of fusion corresponds to the percentage of cells displaying increased cleaved CCF2/AM fluorescence (447 nm).

#### - HIV-1 Infections

After treatment for 18 h, human primary macrophages were infected with HIV-1_ADA_ in 6-well trays with a multiplicity of infection of 0.2, incubated for 24 h at 37°C, and washed with culture medium without FCS. Cells were cultured for 24 h at 37°C in complete culture medium supplemented with library compounds. After 24 h, supernatant was harvested, and cells were lysed for 15 min at 4°C in lysis buffer (20 mM Tris HCl, pH 7.5, 150 mM NaCl, 0.5% NP-40, 50 mM NaF, and 1 mM sodium orthovanadate, supplemented with complete protease inhibitor cocktail; Roche Diagnostic). Lysates were centrifuged at 10,000 *g* for 10 min at 4°C, and the quantity of HIV-1 p24 in the post-nuclear supernatants was determined by ELISA.

## Acknowledgments

The authors gratefully acknowledge the Cell and Tissue Imaging Facility (PICT-IBiSA), Institut Curie, a member of the French National Research Infrastructure, France-BioImaging (ANR10-INBS-04). This work was supported by grants from CNRS, Inserm, Université Paris Descartes, Agence Nationale de la Recherche (2011 BSV3 025 02) and Agence Nationale de Recherches sur le Sida et les Hépatites (ANRS, AO2012-2) to AB, FP and FN, and Fondation pour la Recherche Médicale (FRM DEQ20130326518) to FN. This work has received support under the ‘Investissements d’Avenir’ program launched by the French Government and implemented by ANR with the references ANR-10-LABX-62-IBEID and ANR-10-IDEX-0001-02 PSL. We thank Dr Anna Mularski for careful reading of the manuscript.

## Author contributions

G.B., A.B., F.N. and F.P. designed the study and analyzed data. G.B., F.H. I.S., Y. F. and M.F. performed experiments and analyzed data. S.T., A.L., E.D.N. set-up, performed and analyzed the chemical screening. P.G. conducted the bioinformatics analysis of the screens. G.B., F.H., A.B., F.N., F.H. and F.P. wrote the manuscript, which all coauthors commented on.

## References

1. Alkhatib, G. et al. CC CKR5: a RANTES, MIP-1alpha, MIP-1beta receptor as a fusion cofactor for macrophage-tropic HIV-1. Science 272, 1955–1958 (1996).

2. Choe, H. et al. The beta-chemokine receptors CCR3 and CCR5 facilitate infection by primary HIV-1 isolates. Cell 85, 1135–1148 (1996).

3. Deng, H. et al. Identification of a major co-receptor for primary isolates of HIV-1. Nature 381, 661–666 (1996).

4. Doranz, B.J. et al. A dual-tropic primary HIV-1 isolate that uses fusin and the beta-chemokine receptors CKR-5, CKR-3, and CKR-2b as fusion cofactors. Cell 85, 1149–1158 (1996).

5. Dragic, T. et al. HIV-1 entry into CD4+ cells is mediated by the chemokine receptor CC-CKR-5. Nature 381, 667–673 (1996).

6. Liu, R. et al. Homozygous defect in HIV-1 coreceptor accounts for resistance of some multiply-exposed individuals to HIV-1 infection. Cell 86, 367–377 (1996).

7. Samson, M. et al. Resistance to HIV-1 infection in caucasian individuals bearing mutant alleles of the CCR-5 chemokine receptor gene. Nature 382, 722–725 (1996).

8. Hutter, G. et al. Long-term control of HIV by CCR5 Delta32/Delta32 stem-cell transplantation. N Engl J Med 360, 692–698 (2009).

9. Dorr, P. et al. Maraviroc (UK-427,857), a potent, orally bioavailable, and selective small-molecule inhibitor of chemokine receptor CCR5 with broad-spectrum anti-human immunodeficiency virus type 1 activity. Antimicrob Agents Chemother 49, 4721–4732 (2005).

10. Fatkenheuer, G. et al. Efficacy of short-term monotherapy with maraviroc, a new CCR5 antagonist, in patients infected with HIV-1. Nat Med 11, 1170–1172 (2005).

11. Jacobson, J.M. et al. Antiviral activity of single-dose PRO 140, a CCR5 monoclonal antibody, in HIV-infected adults. J Infect Dis 198, 1345–1352 (2008).

12. Trkola, A. et al. Potent, broad-spectrum inhibition of human immunodeficiency virus type 1 by the CCR5 monoclonal antibody PRO 140. J Virol 75, 579–588 (2001).

13. Perez, E.E. et al. Establishment of HIV-1 resistance in CD4+ T cells by genome editing using zinc-finger nucleases. Nat Biotechnol 26, 808–816 (2008).

14. Tebas, P. et al. Gene editing of CCR5 in autologous CD4 T cells of persons infected with HIV. N Engl J Med 370, 901–910 (2014).

15. Boncompain, G. & Perez, F. The many routes of Golgi-dependent trafficking. Histochem Cell Biol 140, 251–260 (2013).

16. Boncompain, G. et al. Synchronization of secretory protein traffic in populations of cells. Nat Methods 9, 493–498 (2012).

17. Fourriere, L., Divoux, S., Roceri, M., Perez, F. & Boncompain, G. Microtubule-independent secretion requires functional maturation of Golgi elements. J Cell Sci 129, 3238–3250 (2016).

18. Blanpain, C. et al. Palmitoylation of CCR5 is critical for receptor trafficking and efficient activation of intracellular signaling pathways. J Biol Chem 276, 23795–23804 (2001).

19. Percherancier, Y. et al. Palmitoylation-dependent control of degradation, life span, and membrane expression of the CCR5 receptor. J Biol Chem 276, 31936–31944 (2001).

20. Cavrois, M., De Noronha, C. & Greene, W.C. A sensitive and specific enzyme-based assay detecting HIV-1 virion fusion in primary T lymphocytes. Nat Biotechnol 20, 1151–1154 (2002).

21. Benkirane, M., Jin, D.Y., Chun, R.F., Koup, R.A. & Jeang, K.T. Mechanism of transdominant inhibition of CCR5-mediated HIV-1 infection by ccr5delta32. J Biol Chem 272, 30603–30606 (1997).

22. Glass, W.G. et al. CCR5 deficiency increases risk of symptomatic West Nile virus infection. J Exp Med 203, 35–40 (2006).

23. Lim, J.K. et al. Genetic deficiency of chemokine receptor CCR5 is a strong risk factor for symptomatic West Nile virus infection: a meta-analysis of 4 cohorts in the US epidemic. J Infect Dis 197, 262–265 (2008).

24. Kindberg, E. et al. A deletion in the chemokine receptor 5 (CCR5) gene is associated with tickborne encephalitis. J Infect Dis 197, 266–269 (2008).

25. Brass, A.L. et al. Identification of host proteins required for HIV infection through a functional genomic screen. Science 319, 921–926 (2008).

26. Konig, R. et al. Global analysis of host-pathogen interactions that regulate early-stage HIV-1 replication. Cell 135, 49–60 (2008).

27. Park, R.J. et al. A genome-wide CRISPR screen identifies a restricted set of HIV host dependency factors. Nat Genet 49, 193–203 (2017).

28. Zhou, H. et al. Genome-scale RNAi screen for host factors required for HIV replication. Cell Host Microbe 4, 495–504 (2008).

29. Charette, N., Holland, P., Frazer, J., Allen, H. & Dupre, D.J. Dependence on different Rab GTPases for the trafficking of CXCR4 and CCR5 homo or heterodimers between the endoplasmic reticulum and plasma membrane in Jurkat cells. Cell Signal 23, 1738–1749 (2011).

30. Li, C. et al. The GTPase Rab43 Controls the Anterograde ER-Golgi Trafficking and Sorting of GPCRs. Cell Rep 21, 1089–1101 (2017).

31. Fukata, M., Fukata, Y., Adesnik, H., Nicoll, R.A. & Bredt, D.S. Identification of PSD-95 palmitoylating enzymes. Neuron 44, 987–996 (2004).

32. Gottlieb, C.D., Zhang, S. & Linder, M.E. The Cysteine-rich Domain of the DHHC3 Palmitoyltransferase Is Palmitoylated and Contains Tightly Bound Zinc. J Biol Chem 290, 29259–29269 (2015).

33. Tang, L., Qiu, R., Tang, Y. & Wang, S. Cadmium-zinc exchange and their binary relationship in the structure of Zn-related proteins: a mini review. Metallomics 6, 1313–1323 (2014).

34. Tomita, F. & Tamaoki, T. Tetrocarcins, novel antitumor antibiotics. I. Producing organism, fermentation and antimicrobial activity. J Antibiot (Tokyo) 33, 940–945 (1980).

35. Nakashima, T., Miura, M. & Hara, M. Tetrocarcin A inhibits mitochondrial functions of Bcl-2 and suppresses its anti-apoptotic activity. Cancer Res 60, 1229–1235 (2000).

36. Anether, G., Tinhofer, I., Senfter, M. & Greil, R. Tetrocarcin-A--induced ER stress mediates apoptosis in B-CLL cells via a Bcl-2--independent pathway. Blood 101, 4561–4568 (2003).

37. Westby, M. et al. Reduced maximal inhibition in phenotypic susceptibility assays indicates that viral strains resistant to the CCR5 antagonist maraviroc utilize inhibitor-bound receptor for entry. J Virol 81, 2359–2371 (2007).

38. Roche, M. et al. HIV-1 escape from the CCR5 antagonist maraviroc associated with an altered and less-efficient mechanism of gp120-CCR5 engagement that attenuates macrophage tropism. J Virol 85, 4330–4342 (2011).

39. Tilton, J.C. et al. A maraviroc-resistant HIV-1 with narrow cross-resistance to other CCR5 antagonists depends on both N-terminal and extracellular loop domains of drug-bound CCR5. J Virol 84, 10863–10876 (2010).

40. Mazzolini, J. et al. Inhibition of phagocytosis in HIV-1-infected macrophages relies on Nef-dependent alteration of focal delivery of recycling compartments. Blood 115, 4226–4236 (2010).

41. Jordan, M., Schallhorn, A. & Wurm, F.M. Transfecting mammalian cells: optimization of critical parameters affecting calcium-phosphate precipitate formation. Nucleic Acids Res 24, 596–601 (1996).

42. Pleskoff, O. et al. Identification of a chemokine receptor encoded by human cytomegalovirus as a cofactor for HIV-1 entry. Science 276, 1874–1878 (1997).

43. Clavel, F. & Charneau, P. Fusion from without directed by human immunodeficiency virus particles. J Virol 68, 1179–1185 (1994).

